# A caudate-centered network for alexithymia derived from patients with brain tumors

**DOI:** 10.64898/2026.11.06.764956

**Authors:** James Tanner McMahon, Morgan Rose Healey, Yaser Sanchez-Gama, Fabio Campanella, Miran Skrap, Michael Adam Ferguson

**Author notes:** Corresponding author: James Tanner McMahon, Department of Neurosurgery, Mass General Brigham, Harvard Medical School, Boston, MA 02115, USA. These authors contributed equally (James Tanner McMahon, Morgan Rose Healey).

## Abstract

Alexithymia is a trait characterized by impaired emotional awareness. Previous accounts examining the neurobiological underpinnings of this trait have centered on a variety of structures, and alexithymia-causing lesions occur in disparate locations. We use lesion network mapping to derive a neural circuit for alexithymia.

In a neurosurgical cohort of 45 patients with gliomas (19 female; aged 18–75 years), we compute each lesion’s connectivity to the rest of the brain using a normative connectome (n = 1,000) and relate these connectivity maps to preoperative Toronto Alexithymia Scale scores using permutation testing.

Lesions associated with higher alexithymia are connected to a common network that includes the bilateral caudate heads (right > left) and the anterior cingulate cortex, peaking in the right caudate head (family-wise error-corrected p < 0.05). The network converges with coordinates from an independent meta-analysis of gray matter volume in alexithymia (13 of 14 coordinates positive; p = 0.0004). Of the three subscales, difficulty identifying feelings localized to this network. Within the striatum, the network spans the executive and limbic subdivisions.

These findings suggest a caudate-centered circuit, at the interface of affective and executive systems, that unifies disparate lesion locations.

## Introduction

Alexithymia, literally “no words for emotions,” is a trait characterized by difficulty interpreting and expressing one’s own emotional states and by an externally oriented mode of thinking.^1,2^ It varies continuously as a subclinical phenomenon across the general population, is elevated in a range of psychiatric and medical conditions, and can be acquired following brain injury, including traumatic brain injury and stroke.^3,4^ Higher alexithymia is associated with maladaptive coping under emotional stress, elevated risk of medical and psychiatric illness, and poorer health-related quality of life and life satisfaction.^3,5^ Despite this burden, the mechanisms by which emotional experience reaches awareness remain poorly understood, and proposed accounts span developmental, linguistic, cognitive, and neurobiological levels without consensus.^6–10^

Neuroimaging studies have implicated a broad set of regions in alexithymia,^11^ and meta-analyses have identified partial convergence on the amygdala, insula, and dorsal anterior cingulate in functional neuroimaging studies^12^ and on the insula, amygdala, orbitofrontal cortex, and striatum in structural neuroimaging studies.^13^ However, individual findings remain heterogeneous, and because these studies are correlational, they cannot distinguish regions that contribute to alexithymia from those that reflect its consequence or correlates. In a recent advance, Zhang et al.^14^ mapped coordinates from both structural and task-based studies onto a normative connectome, situating alexithymia within large-scale brain networks. This approach, however, drew on a heterogeneous sample from the literature, inherited the correlational limitations of its source studies, and summarized results primarily as overlap with canonical networks.

Lesion studies offer a stronger causal inference, yet implicate disparate sites: the anterior insula,^15^ right anterior cingulate cortex,^16^ left frontoinsular cortex,^17^ anterior and medial prefrontal cortex,^18^ and the right hemisphere more broadly.^19^ Whether these heterogeneous lesions share underlying circuitry remains unknown.

Lesion network mapping (LNM) addresses this question, offering a powerful framework for investigating how disparate lesions converge on a uniform brain network by shifting the focus from lesion location to lesion functional connectivity.^20,21^ This approach effectively addresses the paradox of localization, wherein similar symptoms emerge from neuroanatomically diverse lesions. LNM has been used to identify the neural underpinnings of neuropsychiatric conditions, including depression and post-traumatic stress disorder (PTSD).^22,23^ Further, LNM has also been applied to symptoms arising from brain tumors,^24,25^ supporting its use in neuro-oncological populations.

In this study, we applied LNM to a cohort of patients with intrinsic glioma^18^ to identify brain regions whose connectivity to lesion sites is associated with greater preoperative alexithymia, and to test whether these lesions map onto a common neural network. A previous voxel-based lesion-symptom mapping analysis of this cohort associated alexithymia with anterior and medial prefrontal lesions^18^; here we asked whether lesions across locations converge on a shared network defined by their connectivity. We then tested the convergence of this network with coordinates from a coordinate-based meta-analysis of gray matter volume in alexithymia,^13^ and characterized its anatomy in terms of key structures. Finally, we examined the three components of alexithymia—difficulty identifying feelings (DIF), difficulty describing feelings (DDF), and externally oriented thinking (EOT)—mapping and characterizing the anatomy of significant findings.

## Materials and methods

### Generating lesion network maps

We analyzed patients with glioma from a previously published cohort^18^ who completed the Toronto Alexithymia Scale (TAS-20) before tumor resection; patients with meningioma or metastasis were excluded. Of 50 glioma patients in the original cohort, five were excluded (four because lesion imaging was unavailable and one because full TAS-20 data were unavailable), leaving 45 patients (19 female; age 18–75 years, mean 46.4 ± 15.8; education mean 12.2 ± 3.9 years; WHO grade II, n = 20; III, n = 7; IV, n = 18). The primary outcome was preoperative score in the TAS-20, which assesses difficulty identifying and describing feelings as well as externally oriented thinking.^26,27^ Normality of TAS-20 total and subscale scores was assessed with Shapiro–Wilk tests. Details of recruitment, surgical procedures, and psychometric assessment are provided in the original report.^18^ The original study was approved by the ethical committee of A.O.U. S. Maria della Misericordia of Udine, and all patients gave written informed consent.^18^

Lesion network mapping was performed in line with previously established methods,^21,28^ using the binarized lesion masks generated for voxel-based lesion-symptom mapping by Campanella et al.^18^ (Fig. 1), which were traced on each patient’s preoperative MRI and spatially normalized to MNI space using SPM8, and analyzed on the 2-mm MNI152NLin6Asym template used for lesion network mapping. Postoperative scans were not available for all patients and are confounded by both size of resection and brain deformation in the immediate post-operative period. Lesion masks were used as seeds for resting-state functional connectivity analysis computed against a normative connectome dataset (n = 1,000 healthy right-handed participants; 42.7% male; ages 18–35 years; mean age 21.3 years).^29^ Voxelwise Pearson correlations between each binarized lesion seed and every brain voxel were calculated and Fisher z-transformed, then combined across connectome subjects into a t-statistic map, to produce individual lesion-network maps for each of the 45 patients (Fig. 2).

**Figure 1.**
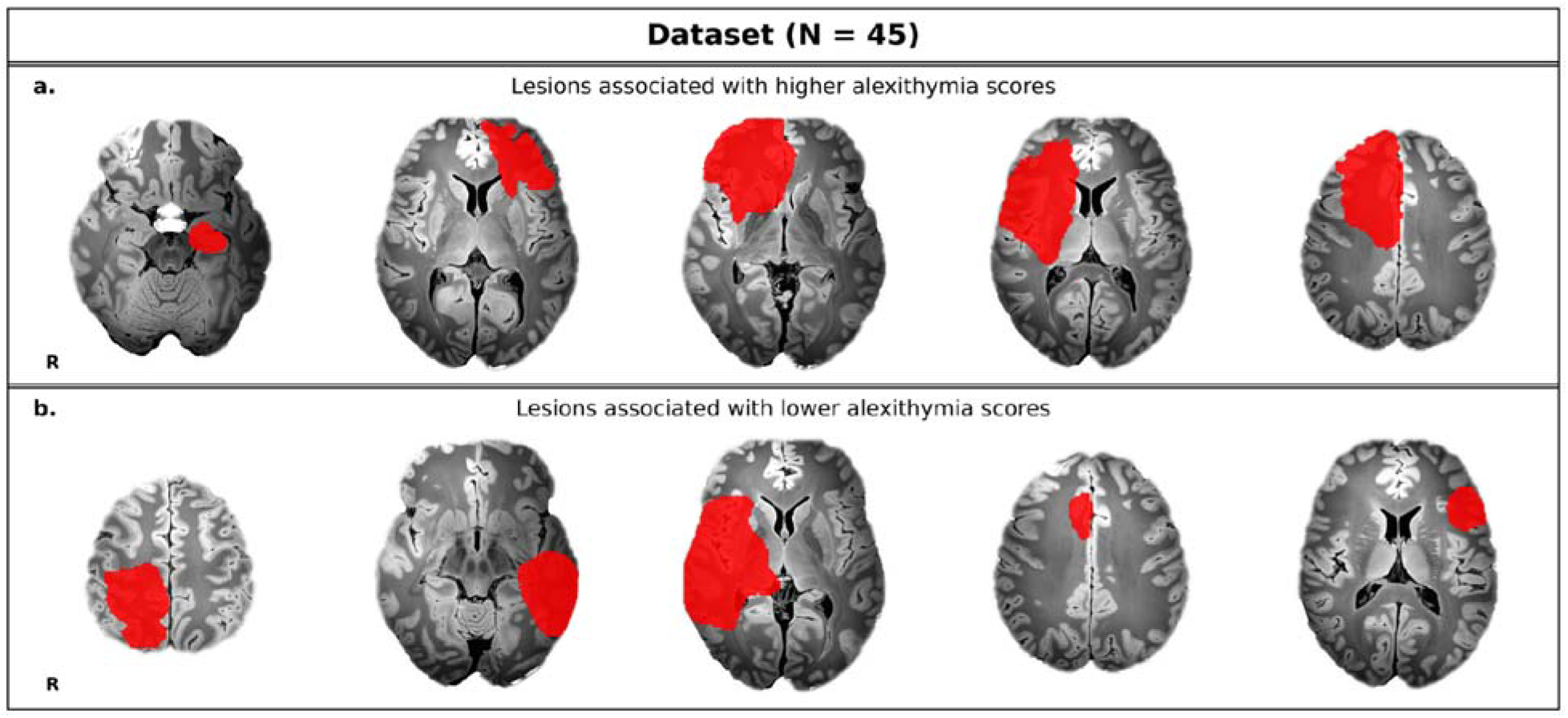
Lesions associated with alexithymia occur in heterogeneous locations. Representative tumor lesion masks (red) from patients with (a) higher and (b) lower preoperative TAS-20 scores, shown on axial slices of the 7 T synthesized FLASH (200 µm) template in MNI space.^33^ Images are in radiological convention (R = right).

**Figure 2.**
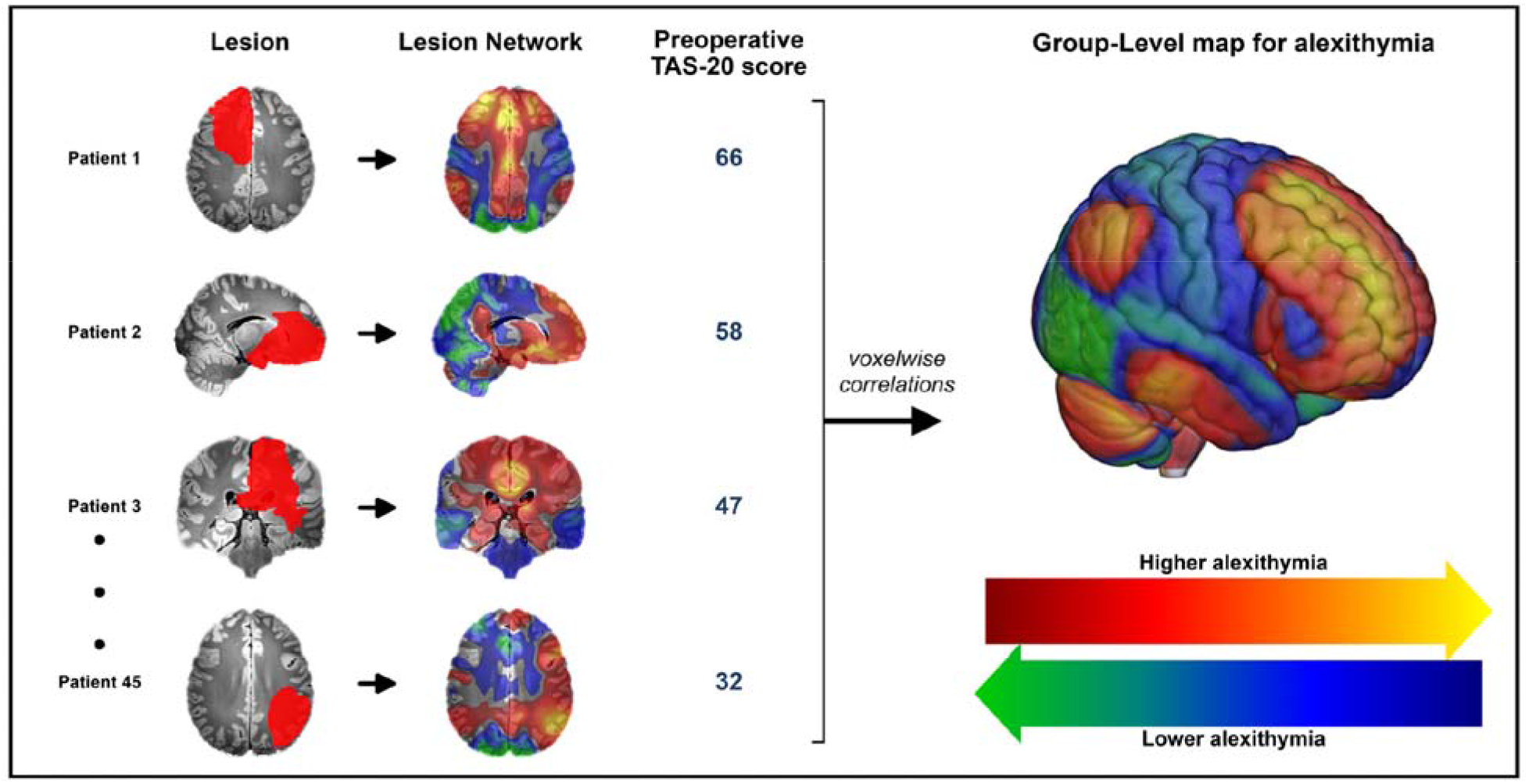
Data-driven method for identifying a lesion network for alexithymia. For each patient, the lesion mask was used as a seed in a normative connectome (n = 1,000) to generate a lesion network map. Lesion network maps from all 45 patients were then related voxelwise to preoperative TAS-20 scores, controlling for age, sex, education and lesion volume, to produce a group-level map. Warm colors indicate voxels where stronger connectivity with lesion sites was associated with higher alexithymia; cool colors, lower alexithymia.

### Deriving the network for alexithymia

To identify a distributed brain network whose functional connectivity with lesion sites corresponds to preoperative alexithymia, we fitted a general linear model, at each voxel, between lesion-seed connectivity and individual preoperative TAS-20 scores, controlling for age, sex, education and lesion volume (continuous variables mean-centered). As such, each voxel received a statistic reflecting the strength of association between connectivity at that location and preoperative TAS-20 score across the 45 patients. Permutation testing with the Freedman–Lane procedure^30^ (FSL randomise; 10,000 permutations)^31^ was used to generate whole-brain t-maps in which positive values indicate that stronger functional connectivity with lesion sites corresponds to higher alexithymia, and negative values to lower alexithymia (Fig. 2). Significance was assessed with two-tailed tests, using voxelwise family-wise error (FWE) and false discovery rate (FDR) correction at p < 0.05 within a dilated MNI152 2 mm brain mask (292,019 voxels). Because our question concerned lesion connectivity associated with greater alexithymia, only positive associations were interpreted. Resulting t-statistic maps were visualized in MRIcroGL (v1.2)^32^ using the 7 T synthesized FLASH (200 µm) template in MNI space.^33^ Results were masked to the brain template for display.

### Convergence using meta-analysis of brain morphometric studies in alexithymia

To identify neuroanatomical commonalities and differences between the lesion-derived alexithymia network and independent structural findings, we obtained coordinates from a coordinate-based meta-analysis of gray matter volume in alexithymia, restricted to whole-brain studies.^13^ These coordinates mark regions of reduced gray matter volume with higher alexithymia. We sampled the unthresholded alexithymia t-map at the nearest voxel to each coordinate and tested whether the mean t-value across coordinates differed from zero using a two-sided one-sample t-test. Positive values indicate that lesions more functionally connected to these regions were associated with higher alexithymia; a mean significantly greater than zero would therefore indicate convergence between the lesion-derived network and regions of structural alteration in alexithymia.

### Characterizing the whole-brain peak

After identifying FWE-significant regions within our network, we characterized the peak region using a published connectivity-based anatomical parcellation of that structure, superimposing each subdivision on the alexithymia map.^34^ We calculated the number of FWE-significant voxels and the mean t-value within each subdivision, and present these as descriptive statistics. Higher mean t-values indicate a stronger association between lesion connectivity and alexithymia within that subdivision.

### Subscale analyses

To test whether any individual TAS-20 subscale was driving the analysis, we repeated the process of fitting a general linear model, at each voxel, between lesion-seed connectivity and individual preoperative TAS-20 subscale scores: difficulty identifying feelings (DIF), difficulty describing feelings (DDF), and externally oriented thinking (EOT). The same covariates and permutation procedure used in the primary analysis were used for the subscale analyses. To account for testing three subscales, results were considered significant at FWE-corrected p < 0.017 (0.05/3).

### Statistical analysis

All tests used α = 0.05; the tests, correction methods, and tails for each analysis are described above. Permutation inference assumes exchangeability of errors under the null hypothesis rather than normally distributed errors. Non-imaging statistics were computed in Python. No a priori sample size calculation was performed because the sample was determined by the available cohort. Blinding and randomization were not applicable to this retrospective observational analysis.

## Results

### Deriving the network for alexithymia

Lesion network mapping across 45 neurosurgical patients revealed a distributed, whole-brain pattern of functional connectivity associated with preoperative alexithymia scores (Fig. 3). Preoperative TAS-20 scores were approximately normally distributed (Shapiro–Wilk W = 0.98, p = 0.58; Supplementary Fig. 1). TAS-20 subscale scores were approximately normally distributed (Shapiro–Wilk p = 0.104, 0.551 and 0.495 for DIF, DDF and EOT, respectively) and showed adequate variability: DIF, 14.3 ± 5.0 (range 7–25); DDF, 12.3 ± 3.6 (range 5–21); EOT, 18.6 ± 3.6 (range 10–26). The resulting network encompassed the dorsal prefrontal cortex, bilateral heads of the caudate, and the bilateral cingulate. The peak voxel was located in the right caudate head (t = 5.63; MNI [14, 22, 4]). The results survived multiple comparison corrections at both FDR and FWE levels. The FDR-thresholded map involved the caudate, anterior cingulate cortex, right frontal pole, right dorsolateral prefrontal cortex, and right thalamus. The FWE map implicated the heads of the caudate bilaterally (right > left) and the anterior cingulate (peak FWE p = 0.005).

**Figure 3.**
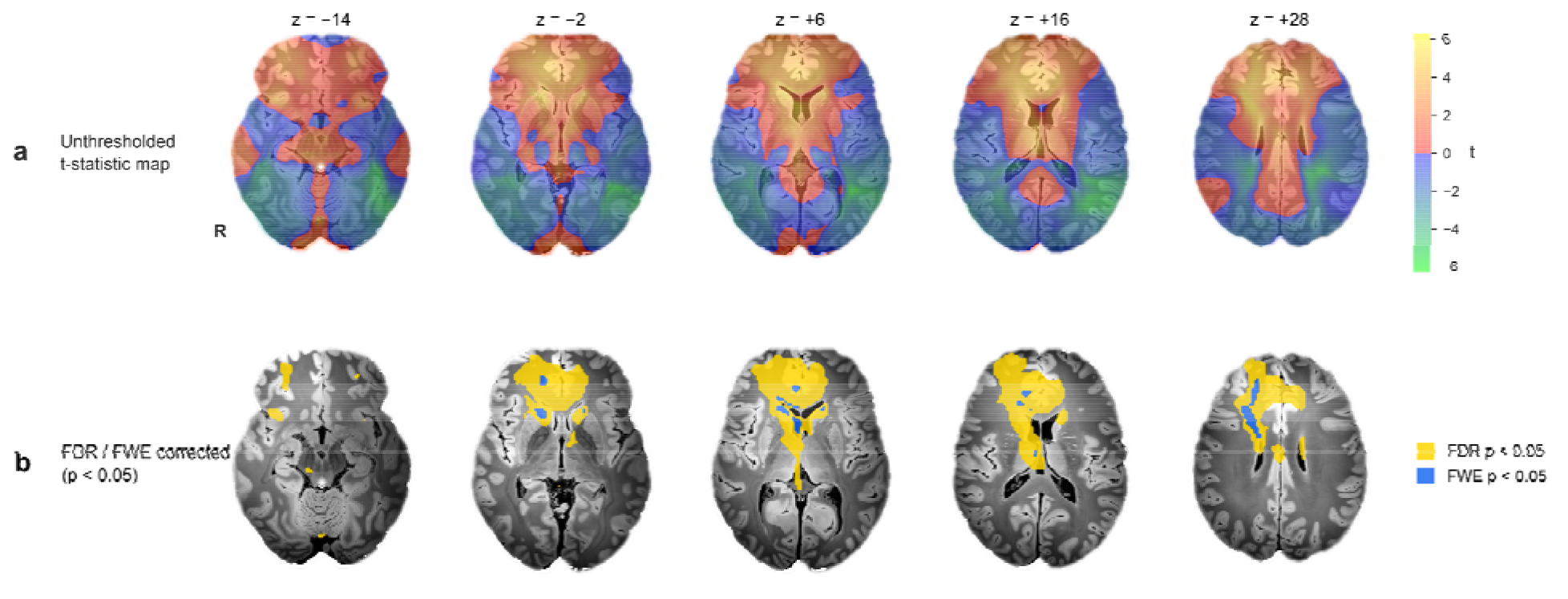
A lesion network for alexithymia. (a) Unthresholded t-statistic map of the association between lesion network connectivity and preoperative TAS-20 score (n = 45), controlling for age, sex, education and lesion volume. Cool colors (negative associations) are shown for completeness and are not interpreted. (b) Voxels surviving correction for multiple comparisons (FSL randomise, Freedman–Lane, 10,000 permutations; two-tailed, positive associations shown): false discovery rate p < 0.05 (yellow) and family-wise error p < 0.05 (blue). The peak lies in the right caudate head (t = 5.63; MNI [14, 22, 4]). Axial slices z = −14, −2, +6, +16, +28; radiological convention.

### Convergence using meta-analysis of brain morphometric studies in alexithymia

Our results were convergent with prior brain morphometric analyses (Table 1). Fourteen coordinates were identified from a meta-analysis of brain morphology in alexithymia.^13^ Sampled at these coordinates, the alexithymia network showed consistently positive values (mean t = 1.77; 13 of 14 coordinates positive; one-sample t(13) = 4.73, p = 0.0004), and four coordinates, all in or adjacent to the right caudate, localized to the FDR-corrected network.

**Table 1.**
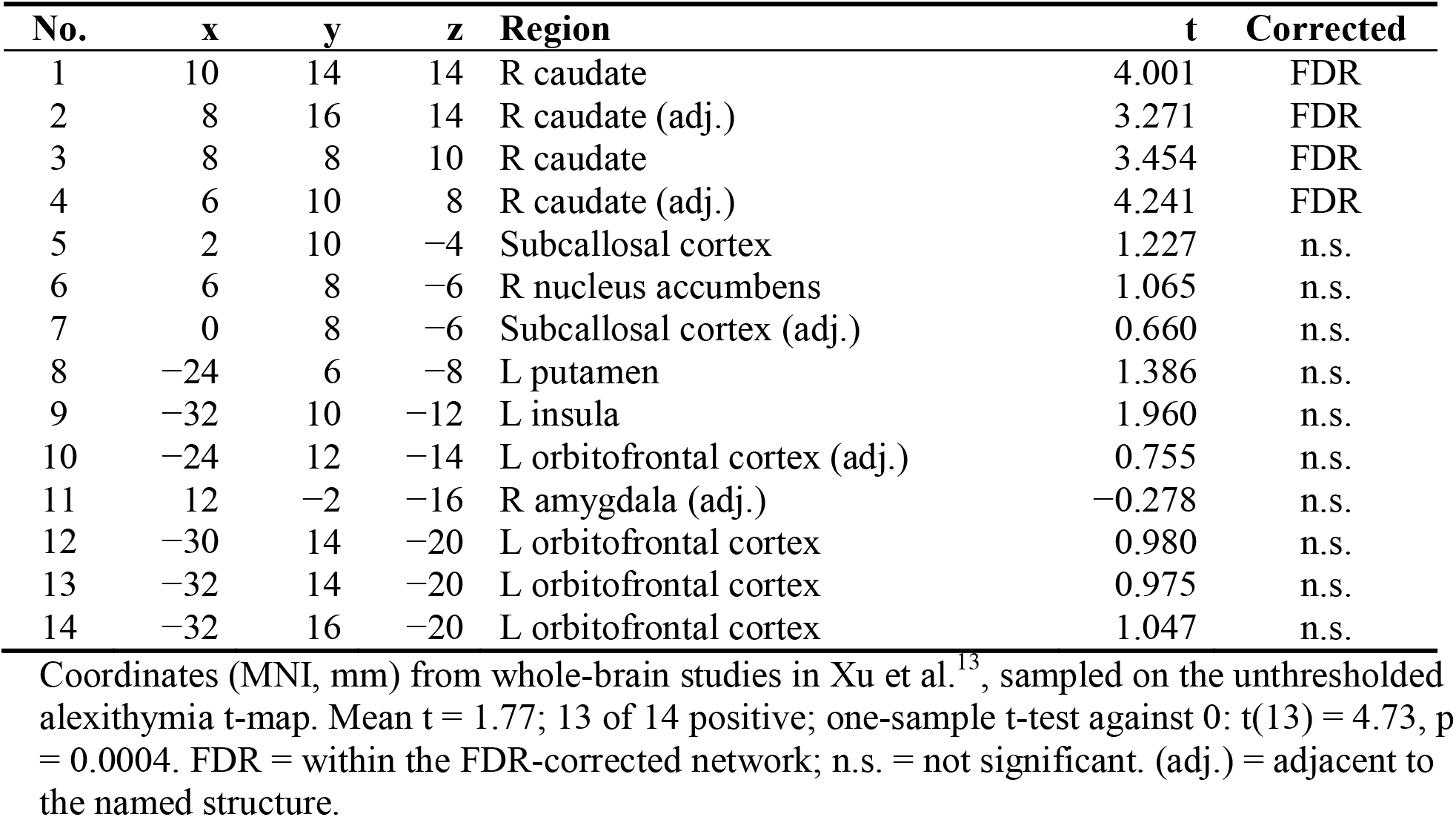
Convergence with a meta-analysis of gray matter volume in alexithymia.

| **No.** | **x** | **y** | **z** | **Region** | **t** | **Corrected** |
|---|---|---|---|---|---|---|
| 1 | 10 | 14 | 14 | R caudate | 4.001 | FDR |
| 2 | 8 | 16 | 14 | R caudate (adj.) | 3.271 | FDR |
| 3 | 8 | 8 | 10 | R caudate | 3.454 | FDR |
| 4 | 6 | 10 | 8 | R caudate (adj.) | 4.241 | FDR |
| 5 | 2 | 10 | −4 | Subcallosal cortex | 1.227 | n.s. |
| 6 | 6 | 8 | −6 | R nucleus accumbens | 1.065 | n.s. |
| 7 | 0 | 8 | −6 | Subcallosal cortex (adj.) | 0.660 | n.s. |
| 8 | −24 | 6 | −8 | L putamen | 1.386 | n.s. |
| 9 | −32 | 10 | −12 | L insula | 1.960 | n.s. |
| 10 | −24 | 12 | −14 | L orbitofrontal cortex (adj.) | 0.755 | n.s. |
| 11 | 12 | −2 | −16 | R amygdala (adj.) | −0.278 | n.s. |
| 12 | −30 | 14 | −20 | L orbitofrontal cortex | 0.980 | n.s. |
| 13 | −32 | 14 | −20 | L orbitofrontal cortex | 0.975 | n.s. |
| 14 | −32 | 16 | −20 | L orbitofrontal cortex | 1.047 | n.s. |

Coordinates (MNI, mm) from whole-brain studies in Xu et al.<sup>13</sup>, sampled on the unthresholded alexithymia t-map. Mean t = 1.77; 13 of 14 positive; one-sample t-test against 0: t(13) = 4.73, p = 0.0004. FDR = within the FDR-corrected network; n.s. = not significant. (adj.) = adjacent to the named structure.

### Characterizing the whole-brain peak

The network peak localized to the caudate, which was therefore selected as our primary region of interest. We characterized it using the Oxford-Imanova Striatal Connectivity Atlas,^34^ which parcellates the striatum into seven subdivisions according to each voxel’s strongest anatomical connectivity with cortical targets, derived from diffusion MRI and probabilistic tractography in 12 healthy adults. The limbic subdivision is defined by connectivity with medial orbitofrontal and subgenual anterior cingulate cortex, and the executive subdivision by connectivity with dorsolateral and frontopolar prefrontal cortex. Of the FWE-significant voxels falling within the striatal subdivisions, the majority (72 of 88) localized to the executive subdivision (Fig. 4; Table 2). A smaller portion localized to the limbic subdivision (16 voxels). No voxels localized to caudal motor, rostral motor, temporal, parietal, or occipital striatal subdivisions. Mean t-values were highest in the executive (t = 2.41) and limbic (t = 1.80) subdivisions (Table 2).

**Figure 4.**
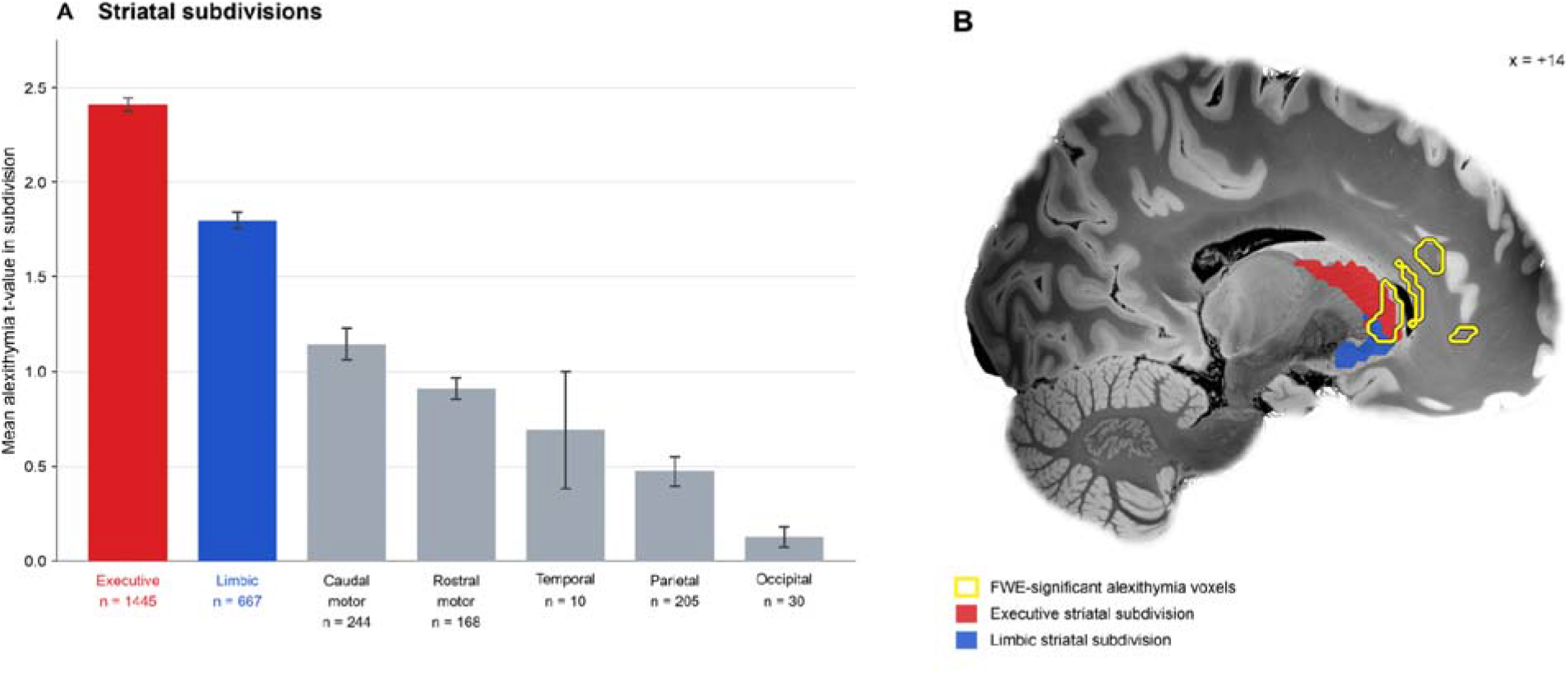
Distribution of the alexithymia network across striatal subdivisions. (A) Mean t-value of the unthresholded alexithymia map within each connectivity-defined striatal subdivision^34^; error bars show SEM across voxels; n = number of 2 mm voxels per subdivision. (B) Sagittal slice (x = +14) showing the executive (red) and limbic (blue) subdivisions and FWE-significant alexithymia voxels (yellow outline).

**Table 2.**
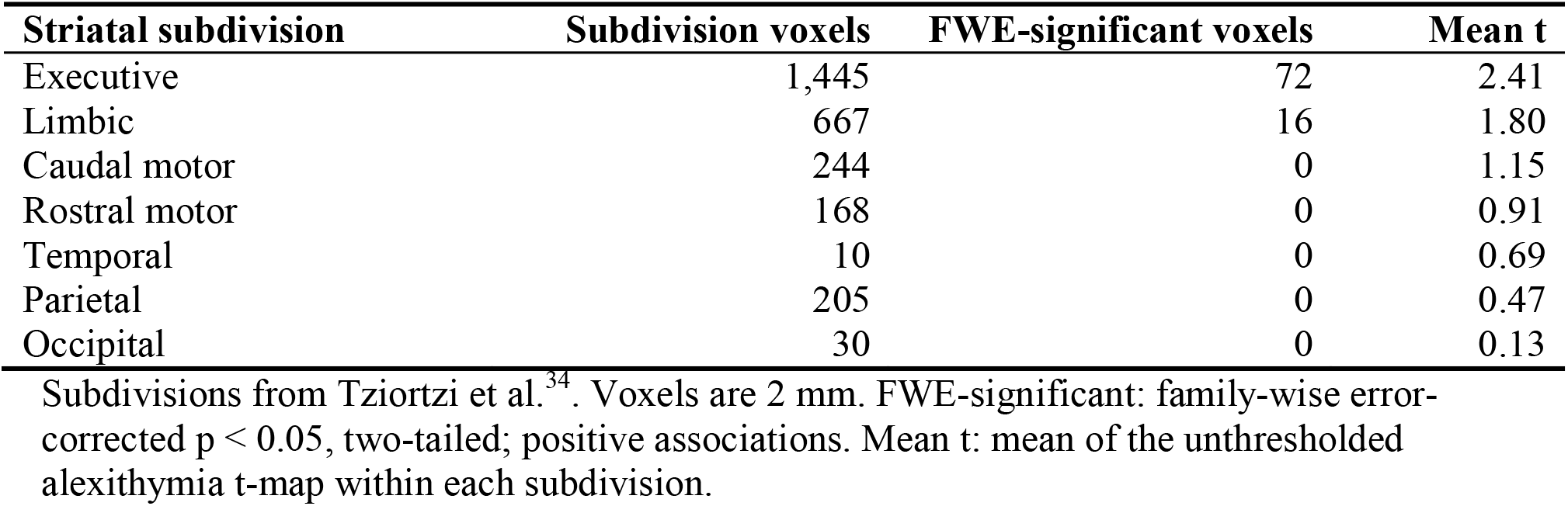
FWE-significant voxels and mean t-values within striatal subdivisions.

### Subscale analyses

Subscale analyses indicated that this association was significant for difficulty identifying feelings (DIF; 238 FWE-significant voxels at p < 0.05, 14,242 FDR-significant voxels) (Fig. 5). The strongest association was similarly observed in the head of the right caudate (t = 5.24; MNI [12, 22, 4]; FWE p = 0.009), which survived the corrected threshold of p < 0.017. No voxels survived correction for difficulty describing feelings (DDF) or externally oriented thinking (EOT).

**Figure 5.**
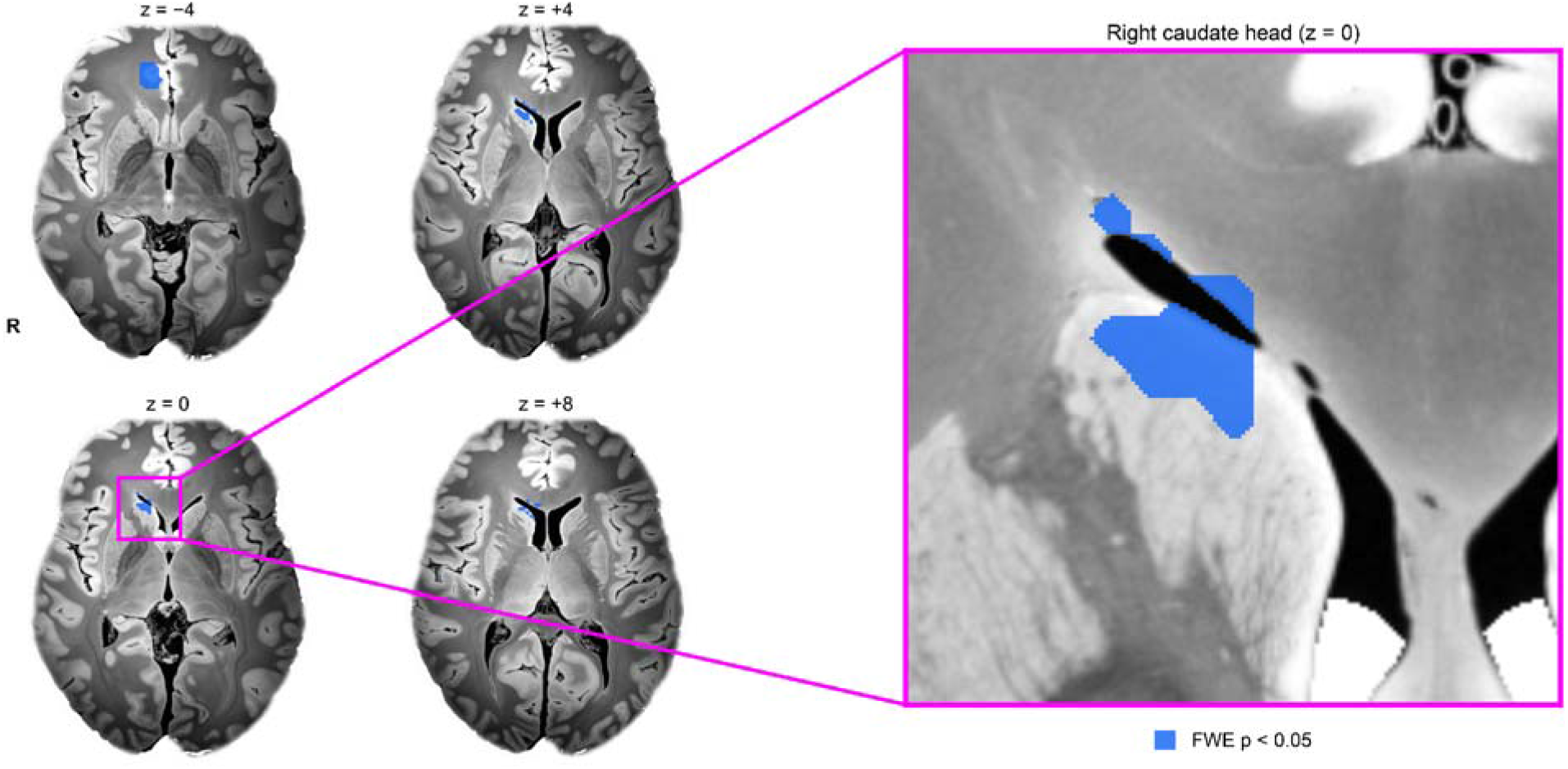
Lesion network for difficulty identifying feelings. Voxels where lesion network connectivity was associated with higher scores on the TAS-20 difficulty identifying feelings subscale (family-wise error p < 0.05, two-tailed, positive associations; FSL randomise, Freedman–Lane, 10,000 permutations; n = 45; same covariates as the primary analysis). The right caudate head survives the subscale-corrected threshold (FWE p < 0.017). Axial slices z = −4, 0, +4, +8; the boxed region at z = 0 is enlarged to show the right caudate head (the cluster peak, t = 5.24, MNI [12, 22, 4], lies at z = +4). A second cluster lies in the pregenual anterior cingulate/paracingulate cortex. For display, the overlay is not shown over cerebrospinal fluid. Radiological convention.

## Discussion

We derive a neural circuit for alexithymia in a cohort of intrinsic glioma patients using lesion network mapping procedures, a technique for localizing clinical symptoms to distributed functional networks.^20^ Our network peaked in the right caudate head, which survived family-wise error (FWE) correction. The false discovery rate (FDR) significant network also includes the bilateral caudate heads (right > left), anterior and paracingulate cortex bilaterally, the right frontal pole, right dorsolateral prefrontal cortex, right thalamus, orbitofrontal cortex and insula.

Previous work has demonstrated alexithymia lesions in disparate locations, including the anterior insula,^15^ right anterior cingulate,^16^ left frontoinsular cortex,^17^ inferior frontal gyrus,^35^ anterior and medial prefrontal cortex,^18^ and the right hemisphere more broadly.^19^ Our results suggest these heterogeneous lesions may share a common circuit: lesions associated with alexithymia were functionally connected to the right caudate head, and several of these previously implicated sites fell within the FDR-corrected network, which was strongly right-lateralized.

Our findings are supported by an independent coordinate-based meta-analysis of gray matter volume in alexithymia,^13^ whose coordinates localized to positive values of our unthresholded network (t(13) = 4.73, p = 0.0004), and four coordinates were encompassed by our FDR-significant network. All four coordinates inside the FDR-significant network were located in or adjacent to the right caudate, implying that lesion-derived and structural findings may point to the same circuit. As such, two independent lines of evidence, lesion connectivity and gray matter morphometry, converge on the right caudate.

In the context of alexithymia, the most notable region of interest has consistently centered on the insula.^15,36,37^ The insula, a core paralimbic region, has repeatedly been linked to interoception and emotional processing and thus is a strong candidate for a core hub that, when disrupted, may contribute to alexithymia.^38,39^ Convergent with this, the right anterior insula fell within the FDR-corrected network. Notably, however, our network-based approach extended the focus from the insula to also include a caudate-centered cortico-striatal network.

Prior work has linked the caudate to emotion.^40–42^ Within alexithymia specifically, prior work has demonstrated reduced caudate or striatal volume with higher alexithymia.^13,43^ Lee et al.^44^ found a negative association between self-reported alexithymia and right caudate activation to emotional stimuli. Similarly, Suslow et al.^45^ observed less right caudate activation in patients with alexithymia than non-alexithymic patients during emotion processing. Our finding of a network peak in the right caudate is consistent with this previous work.

While exploratory, our results revealed that the association with the caudate was concentrated within its executive and limbic subdivisions.^34^ This provides more context for our findings: the limbic zone is linked to medial orbitofrontal and subgenual anterior cingulate cortex, while the executive zone is linked to dorsolateral prefrontal cortex^34^; the former regions in particular have been linked to alexithymia.^13,43^ That the voxels spatially lie on the border of the executive and limbic parcellations is particularly interesting given that the phenotype is characterized by the interaction between the experience of emotion (limbic) and higher-order processing (executive), and that cortical inputs from limbic (orbitofrontal, anterior cingulate) and dorsal prefrontal regions converge within the caudate, providing an anatomical substrate for interaction between affective and executive circuits.^46^ Together, these findings suggest that alexithymia may arise not from damage to a single affective or executive region, but from disruption of their interaction at the caudate.

Of the three TAS-20 subscales, difficulty identifying feelings (DIF) localized significantly to the network. DIF reflects difficulty at the appraisal stage of emotion processing.^9^ Consistent with this, neuroimaging work has linked the cognitive dimension of alexithymia, which includes difficulty identifying feelings, to reduced caudate volume.^43^ As such, our results suggest difficulty recognizing one’s own emotions may localize to caudate-related circuitry.

Altogether, our results support the conceptualization of alexithymia as a network-based trait.^14^ Further, they imply that lesions functionally connected to the caudate contribute to alexithymia, suggesting more attention be given to the caudate when considering neurobiological models of alexithymia. Beyond alexithymia, our results add lesion-based evidence to the caudate’s established non-motor roles,^47^ including emotion^40,42^ and anticipation of reward,^48,49^ by implicating caudate-centered circuitry in emotional awareness. This is consistent with the limbic cortico-subcortical-cortical loop described by Alexander et al.^50^, suggesting that the architecture underlying motor planning and error correction may support emotion processing and regulation in an analogous way.

These results have clinical relevance: alexithymia is a well-recognized transdiagnostic risk factor that predicts less effective psychotherapy,^51^ though it may be modifiable through psychological intervention.^52,53^ That difficulty identifying feelings significantly localized to this network may suggest that interventions directly targeting the appraisal of emotional states, such as emotion-awareness training,^5^ could be particularly relevant. Whether such interventions engage caudate circuitry is a testable question for future work.

There are limitations to our study. We are limited by a modest sample size (n = 45) and the lack of a second dataset for independent validation. The convergence of our network with an independent meta-analysis of brain morphometric studies in alexithymia^13^ partially mitigates this, although it does not substitute for replication. In addition, the convergence test treated meta-analytic coordinates as independent, although several were neighboring peaks of the same cluster (e.g., four in or adjacent to the right caudate), which may overstate its significance. Alexithymia was assessed by self-report, which may be limited in a construct defined by reduced emotional awareness, although the TAS-20 remains the most widely used and validated measure.^26,27^ Lesion network maps were derived from a normative connectome of healthy young adults, which may not reflect connectivity in older patients or in brains altered by tumor. We use preoperative scores, which capture the effect of the tumor before surgical resection.^18^ Our results, as with any lesion mapping study, are limited by the compensatory responses that occur after lesion onset; this may be especially pronounced for slow-growing gliomas, which allow substantial functional reorganization.^54^ However, as such reorganization would be expected to attenuate rather than create lesion-symptom associations, our estimates are likely conservative.

In summary, lesions associated with alexithymia in patients with intrinsic glioma map to a common functional network centered on the right caudate head, a finding that converges with independent morphometric evidence and to which difficulty identifying feelings localizes. To our knowledge, this is the first study to map alexithymia to a common brain network using lesion network mapping. These results extend attention beyond the insula to include caudate-centered cortico-striatal circuitry as a substrate of emotional awareness, and they identify the border between the caudate’s limbic and executive zones as a candidate site where affective experience and its top-down recognition interact. Replication in larger, independent lesion cohorts, including patients with acute lesions such as stroke, will be needed to confirm this circuit and to test whether it can guide more targeted approaches to the assessment and treatment of alexithymia.

## Data availability

De-identified lesion masks and TAS-20 scores are available from F.C. upon reasonable request, subject to approval by the ethical committee of A.O.U. S. Maria della Misericordia of Udine. The normative connectome is available from the Brain Genomics Superstruct Project. Meta-analytic coordinates are reported in Xu et al.^13^ and Table 1. Code used for the analyses and figures is available from the corresponding author upon reasonable request.

## Acknowledgements

We thank the patients who participated in the original study. During the preparation of this work, the authors used Claude (Anthropic) to assist with literature searching and citation verification, writing and debugging analysis and figure-generation code, and language editing and writing assistance. All analyses, code outputs, and cited sources were checked by the authors, who reviewed and edited all content and take full responsibility for the content of the publication.

## Author contributions

J.T.M. and M.R.H. contributed equally to this work. **Conceptualization:** J.T.M., M.A.F. **Methodology:** J.T.M., M.R.H., Y.S.-G., M.A.F. **Formal analysis:** M.R.H., Y.S.-G., J.T.M., M.A.F. **Investigation:** F.C., M.S. **Resources:** F.C., M.S. **Data curation:** F.C., J.T.M., M.R.H. **Visualization:** M.R.H., Y.S.-G. **Project administration:** M.R.H., J.T.M**. Writing – original draft:** J.T.M., M.R.H. **Writing – review & editing:** all authors. **Supervision:** M.A.F.

## Funding

This research received no specific grant from any funding agency in the public, commercial, or not-for-profit sectors.

## Competing interests

The authors report no competing interests.

## Supplementary Figures

**Supplementary Fig. 1.**
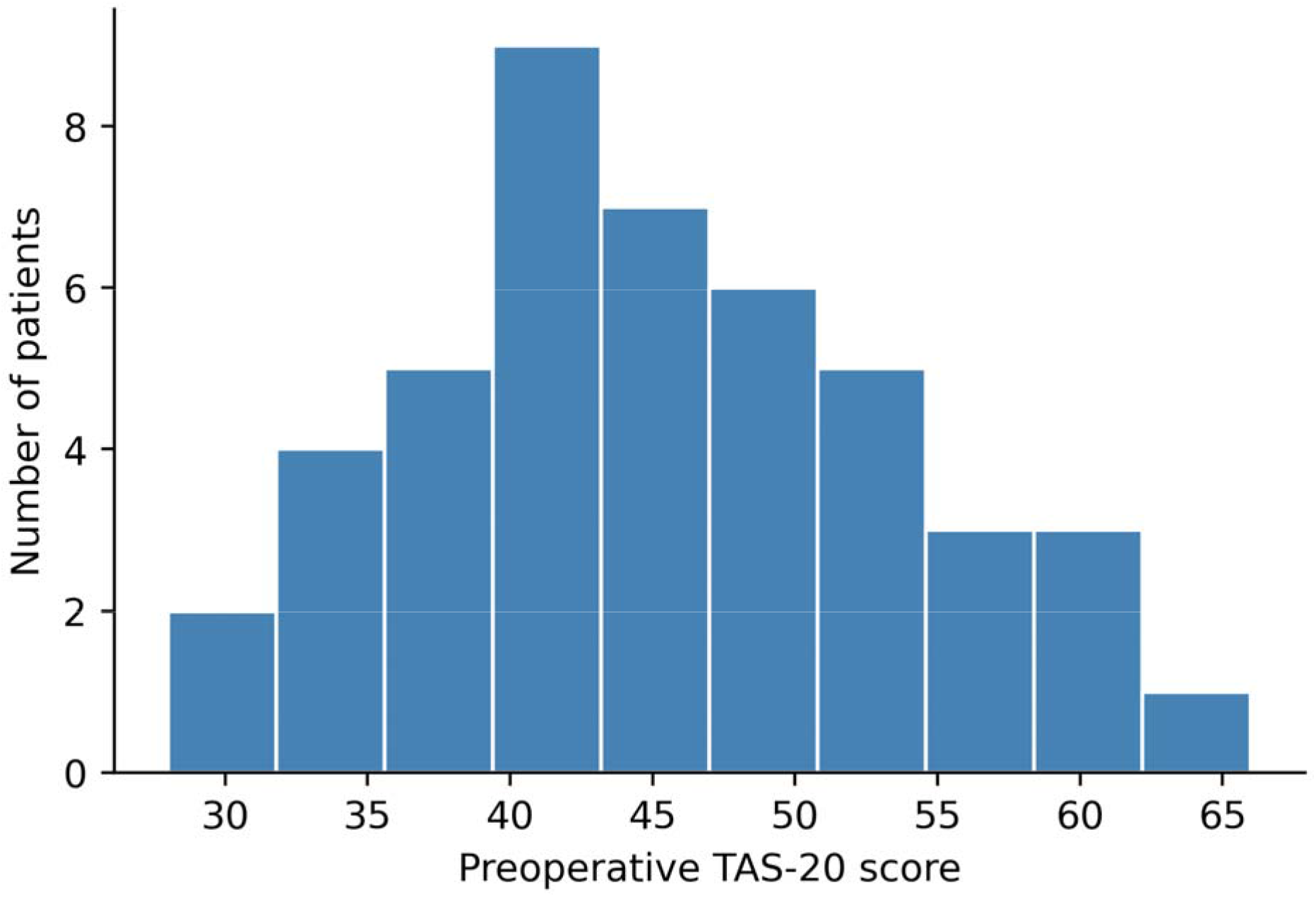
Distribution of preoperative alexithymia scores. Histogram of preoperative Toronto Alexithymia Scale (TAS-20) total scores in the 45 patients included in the lesion network mapping analysis (mean 45.2 ± 8.8; range 28–66). Scores were approximately normally distributed (Shapiro–Wilk W = 0.98, p = 0.58). Three patients scored in the alexithymic range (≥ 61) and eight in the possible-alexithymia range (52–60).

## Notes

### Competing Interest Statement

The authors have declared no competing interest.

## References

1. Sifneos PE. The Prevalence of “Alexithymic” Characteristics in Psychosomatic Patients. Psychother Psychosom. 1973;22(2/6):255–262. doi:10.1159/000286529

2. Taylor G. Alexithymia: concept, measurement, and implications for treatment. Am J Psychiatry. 1984;141(6):725–732. doi:10.1176/ajp.141.6.725

3. Hogeveen J, Grafman J. Alexithymia. Handb Clin Neurol. 2021;183:47–62. doi:10.1016/B978-0-12-822290-4.00004-9

4. Parker JDA, Keefer KV, Taylor GJ, Bagby RM. Latent structure of the alexithymia construct: A taxometric investigation. Psychol Assess. 2008;20(4):385–396. doi:10.1037/a0014262

5. Luminet O, Nielson KA. Alexithymia: Toward an Experimental, Processual Affective Science with Effective Interventions. Annu Rev Psychol. 2025;76(1):741–769. doi:10.1146/annurev-psych-021424-030718

6. Hobson H, Brewer R, Catmur C, Bird G. The Role of Language in Alexithymia: Moving Towards a Multiroute Model of Alexithymia. Emot Rev. 2019;11(3):247–261. doi:10.1177/1754073919838528

7. Lee KS, Murphy J, Catmur C, Bird G, Hobson H. Furthering the language hypothesis of alexithymia: An integrated review and meta-analysis. Neurosci Biobehav Rev. 2022;141:104864. doi:10.1016/j.neubiorev.2022.104864

8. Li R, Kajanoja J, Karlsson L, Karlsson H, Karukivi M. Sex-specific role of alexithymia in associations between parental bonding and mental health: A moderated mediation model. J Clin Psychol. 2023;79(1):126–142. doi:10.1002/jclp.23372

9. Preece D, Becerra R, Allan A, Robinson K, Dandy J. Establishing the theoretical components of alexithymia via factor analysis: Introduction and validation of the attention-appraisal model of alexithymia. Personal Individ Differ. 2017;119:341–352. doi:10.1016/j.paid.2017.08.003

10. Schimmenti A, Caretti V. Attachment, Trauma, and Alexithymia. In: Luminet O, Bagby RM, Taylor GJ, eds. Alexithymia. Cambridge University Press; 2018:127–141. doi:10.1017/9781108241595.010

11. Goerlich KS, Aleman A. Neuroimaging Studies of Alexithymia. In: Luminet O, Bagby RM, Taylor GJ, eds. Alexithymia. Cambridge University Press; 2018:207–249. doi:10.1017/9781108241595.015

12. Van Der Velde J, Servaas MN, Goerlich KS, et al. Neural correlates of alexithymia: A meta-analysis of emotion processing studies. Neurosci Biobehav Rev. 2013;37(8):1774–1785. doi:10.1016/j.neubiorev.2013.07.008

13. Xu P, Opmeer EM, Van Tol MJ, Goerlich KS, Aleman A. Structure of the alexithymic brain: A parametric coordinate-based meta-analysis. Neurosci Biobehav Rev. 2018;87:50–55. doi:10.1016/j.neubiorev.2018.01.004

14. Zhang L, Cui X, Ou Y, Liu Y, Lang B, Guo W. Brain network mapping of structural and functional alterations associated with alexithymia. Transl Psychiatry. 2026;16(1):505. doi:10.1038/s41398-026-04259-0

15. Hogeveen J, Bird G, Chau A, Krueger F, Grafman J. Acquired alexithymia following damage to the anterior insula. Neuropsychologia. 2016;82:142–148. doi:10.1016/j.neuropsychologia.2016.01.021

16. Schäfer R, Popp K, Jörgens S, Lindenberg R, Franz M, Seitz RJ. Alexithymia-like Disorder in Right Anterior Cingulate Infarction. Neurocase. 2007;13(3):201–208. doi:10.1080/13554790701494964

17. Hogeveen J, Krueger F, Grafman J. Association Between Alexithymia and Impaired Reward Valuation in Patients with Fronto-insular Damage. Emotion. 2021;21(1):137–147. doi:10.1037/emo0000676

18. Campanella F, Shallice T, Ius T, Fabbro F, Skrap M. Impact of brain tumour location on emotion and personality: a voxel-based lesion–symptom mapping study on mentalization processes. Brain. 2014;137(9):2532–2545. doi:10.1093/brain/awu183

19. Fricchione G, Howanitz E. Aprosodia and Alexithymia - A Case Report. Psychother Psychosom. 1985;43(3):156–160. doi:10.1159/000287873

20. Boes AD, Prasad S, Liu H, et al. Network localization of neurological symptoms from focal brain lesions. Brain. 2015;138(10):3061–3075. doi:10.1093/brain/awv228

21. Fox MD. Mapping symptoms to brain networks with the human connectome. N Engl J Med. 2018;379(23):2237–2245. doi:10.1056/NEJMra1706158

22. Padmanabhan JL, Cooke D, Joutsa J, et al. A Human Depression Circuit Derived From Focal Brain Lesions. Biol Psychiatry. 2019;86(10):749–758. doi:10.1016/j.biopsych.2019.07.023

23. Siddiqi SH, Philip NS, Palm ST, et al. A potential target for noninvasive neuromodulation of PTSD symptoms derived from focal brain lesions in veterans. Nat Neurosci. 2024;27(11):2231–2239. doi:10.1038/s41593-024-01772-7

24. Li Y, Jin Y, Wu D, Zhang L. A depression network caused by brain tumours. Brain Struct Funct. 2022;227(8):2787–2795. doi:10.1007/s00429-022-02573-z

25. Mansouri AM, Germann J, Boutet A, et al. Identification of neural networks preferentially engaged by epileptogenic mass lesions through lesion network mapping analysis. Sci Rep. 2020;10(1):10989. doi:10.1038/s41598-020-67626-x

26. Bagby RM, Parker JD, Taylor GJ. The twenty-item Toronto Alexithymia Scale—I. Item selection and cross-validation of the factor structure. J Psychosom Res. 1994;38(1):23–32. doi:10.1016/0022-3999(94)90005-1

27. Bagby RM, Taylor GJ, Parker JDA. The twenty-item Toronto Alexithymia Scale—II. Convergent, discriminant, and concurrent validity. J Psychosom Res. 1994;38(1):33–40. doi:10.1016/0022-3999(94)90006-X

28. Ferguson MA, Schaper FLWVJ, Cohen A, et al. A neural circuit for spirituality and religiosity derived from patients with brain lesions. Biol Psychiatry. 2022;91(4):380–388. doi:10.1016/j.biopsych.2021.06.016

29. Yeo BTT, Krienen FM, Sepulcre J, et al. The organization of the human cerebral cortex estimated by intrinsic functional connectivity. J Neurophysiol. 2011;106(3):1125–1165. doi:10.1152/jn.00338.2011

30. Freedman D, Lane D. A Nonstochastic Interpretation of Reported Significance Levels. J Bus Econ Stat. 1983;1(4):292–298. doi:10.1080/07350015.1983.10509354

31. Winkler AM, Ridgway GR, Webster MA, Smith SM, Nichols TE. Permutation inference for the general linear model. NeuroImage. 2014;92:381–397. doi:10.1016/j.neuroimage.2014.01.060

32. Rorden C. MRIcroGL: voxel-based visualization for neuroimaging. Nat Methods. 2025;22(8):1613–1614. doi:10.1038/s41592-025-02763-7

33. Edlow BL, Mareyam A, Horn A, et al. 7 Tesla MRI of the ex vivo human brain at 100 micron resolution. Sci Data. 2019;6(1):244. doi:10.1038/s41597-019-0254-8

34. Tziortzi AC, Haber SN, Searle GE, et al. Connectivity-Based Functional Analysis of Dopamine Release in the Striatum Using Diffusion-Weighted MRI and Positron Emission Tomography. Cereb Cortex. 2014;24(5):1165–1177. doi:10.1093/cercor/bhs397

35. Hobson H, Hogeveen J, Brewer R, et al. Language and alexithymia: Evidence for the role of the inferior frontal gyrus in acquired alexithymia. Neuropsychologia. 2018;111:229–240. doi:10.1016/j.neuropsychologia.2017.12.037

36. Bird G, Silani G, Brindley R, White S, Frith U, Singer T. Empathic brain responses in insula are modulated by levels of alexithymia but not autism. Brain. 2010;133(5):1515–1525. doi:10.1093/brain/awq060

37. Silani G, Bird G, Brindley R, Singer T, Frith C, Frith U. Levels of emotional awareness and autism: An fMRI study. Soc Neurosci. 2008;3(2):97–112. doi:10.1080/17470910701577020

38. Craig AD. How do you feel — now? The anterior insula and human awareness. Nat Rev Neurosci. 2009;10(1):59–70. doi:10.1038/nrn2555

39. Critchley HD, Wiens S, Rotshtein P, Öhman A, Dolan RJ. Neural systems supporting interoceptive awareness. Nat Neurosci. 2004;7(2):189–195. doi:10.1038/nn1176

40. Arsalidou M, Duerden EG, Taylor MJ. The centre of the brain: Topographical model of motor, cognitive, affective, and somatosensory functions of the basal ganglia. Hum Brain Mapp. 2013;34(11):3031–3054. doi:10.1002/hbm.22124

41. Mendez MF, Adams NL, Lewandowski KS. Neurobehavioral changes associated with caudate lesions. Neurology. 1989;39(3):349–354. doi:10.1212/WNL.39.3.349

42. Pierce JE, Péron J. The basal ganglia and the cerebellum in human emotion. Soc Cogn Affect Neurosci. 2020;15(5):599–613. doi:10.1093/scan/nsaa076

43. Goerlich-Dobre KS, Votinov M, Habel U, Pripfl J, Lamm C. Neuroanatomical profiles of alexithymia dimensions and subtypes. Hum Brain Mapp. 2015;36(10):3805–3818. doi:10.1002/hbm.22879

44. Lee BT, Lee HY, Park SA, et al. Neural Substrates of Affective Face Recognition in Alexithymia: A Functional Magnetic Resonance Imaging Study. Neuropsychobiology. 2011;63(2):119–124. doi:10.1159/000318086

45. Suslow T, Kugel H, Rufer M, et al. Alexithymia is associated with attenuated automatic brain response to facial emotion in clinical depression. Prog Neuropsychopharmacol Biol Psychiatry. 2016;65:194–200. doi:10.1016/j.pnpbp.2015.10.006

46. Haber SN, Knutson B. The Reward Circuit: Linking Primate Anatomy and Human Imaging. Neuropsychopharmacology. 2010;35(1):4–26. doi:10.1038/npp.2009.129

47. Grahn JA, Parkinson JA, Owen AM. The cognitive functions of the caudate nucleus. Prog Neurobiol. 2008;86(3):141–155. doi:10.1016/j.pneurobio.2008.09.004

48. Salimpoor VN, Benovoy M, Larcher K, Dagher A, Zatorre RJ. Anatomically distinct dopamine release during anticipation and experience of peak emotion to music. Nat Neurosci. 2011;14(2):257–262. doi:10.1038/nn.2726

49. Kawagoe R, Takikawa Y, Hikosaka O. Expectation of reward modulates cognitive signals in the basal ganglia. Nat Neurosci. 1998;1(5):411–416. doi:10.1038/1625

50. Alexander GE, DeLong MR, Strick PL. Parallel organization of functionally segregated circuits linking basal ganglia and cortex. Annu Rev Neurosci. 1986;9(1):357–381. doi:10.1146/annurev.ne.09.030186.002041

51. Ogrodniczuk JS, Piper WE, Joyce AS. Effect of alexithymia on the process and outcome of psychotherapy: A programmatic review. Psychiatry Res. 2011;190(1):43–48. doi:10.1016/j.psychres.2010.04.026

52. Cameron K, Ogrodniczuk J, Hadjipavlou G. Changes in Alexithymia Following Psychological Intervention: A Review. Harv Rev Psychiatry. 2014;22(3):162–178. doi:10.1097/HRP.0000000000000036

53. Norman H, Marzano L, Coulson M, Oskis A. Effects of mindfulness-based interventions on alexithymia: a systematic review. Evid Based Ment Health. 2019;22(1):36–43. doi:10.1136/ebmental-2018-300029

54. Desmurget M, Bonnetblanc F, Duffau H. Contrasting acute and slow-growing lesions: a new door to brain plasticity. Brain. 2007;130(4):898–914. doi:10.1093/brain/awl300

